# Hedgehog pathway activation through conformational blockade of the Patched sterol conduit

**DOI:** 10.1101/783290

**Authors:** Yunxiao Zhang, Wan-Jin Lu, David P. Bulkley, Jiahao Liang, Arthur Ralko, Kelsey J. Roberts, Anping Li, Wonhwa Cho, Yifan Cheng, Aashish Manglik, Philip A. Beachy

## Abstract

Activation of the Hedgehog pathway may have therapeutic value for improved bone healing, taste receptor cell regeneration, and alleviation of colitis or other conditions. Systemic pathway activation, however, may be detrimental and therapeutic application has been difficult for lack of agents amenable to tissue targeting. We have developed a novel agonist, a conformation-specific nanobody against the Hedgehog receptor Patched1. This nanobody potently activates the Hedgehog pathway *in vitro* and *in vivo* by stabilizing an alternative conformation of a Patched1 “switch helix”, as revealed by cryo-EM structure determination. Although this conformation likely constitutes part of the transport cycle, nanobody-trapping disrupts the cycle and prevents substrate movement through the Patched1 sterol conduit. Our conformation-selective nanobody approach provides a new route to the development of transporter-related pharmacologic agents and may be generally applicable to the study of other transporters.

---

Therapeutic manipulation of the Hedgehog pathway has found its clearest application in patients with malignancies whose initiation and growth depend on pathway-activating mutations in the primary cells of the tumor, such as medulloblastoma and basal cell carcinoma ^1, 2^. Pathway blockade, imposed by systemically administered small molecules, has provided clear benefits in such patients, albeit accompanied by unwanted but largely tolerable class effects such as muscle cramps, loss of hair, and loss of taste sensation ^3^. In contrast to promoting tumor growth, however, pathway activity recently has been found to suppress cancer growth and progression when it occurs in stromal cells rather than primary cells ^4^, particularly in cancers of endodermal organs, such as bladder carcinoma, and colon and pancreatic adenocarcinoma ^4–9^. Pathway activation may also confer therapeutic benefits in regeneration of taste receptor cells of the tongue ^10^, which are often lost or diminished in chemotherapy patients, in protection or recovery from diseases such as colitis ^9^, reduction of tissue overgrowth in prostatic hypertrophy ^11^, or acceleration of bone healing in diabetes ^12^.

Despite the potential benefits of pathway activation and the availability of potent small molecule pathway activators ^13^, however, systemic pathway-activating therapies have not been pursued clinically, in part because such treatments are likely to cause overgrowth of mesenchyme and potential initiation or exacerbation of fibrosis in many organs ^6, 14^. These dangerous side effects might be avoided by restricting pathway activation to specific organs or tissue compartments. Restricted pathway activation could be accomplished through conjugation of a pathway agonist to another agent, such as an antibody, that targets specific tissues or organs. Such targeting would be most easily accomplished with a synthetic or genetically encoded peptide as the agonist. Use of the Hedgehog protein for this purpose, however, is hampered by the requirement of lipid modifications for its activity ^15, 16^; these modifications include a cholesteryl moiety on its carboxy-terminus, acquired during auto-processing of the Hedgehog precursor ^17^, and a palmitoyl adduct on its amino-terminus, catalyzed by the acyltransferase Ski1/HHAT ^18^. Other synthetic or genetically-encoded peptides that could easily be conjugated to targeting agents are currently lacking.

Recent insights into the mechanism of Hedgehog signal transduction suggest a route to remedying this deficiency. The primary receptor for Hedgehog is Patched1 (PTCH1), which maintains pathway quiescence by suppressing Smoothened (SMO) a downstream G-protein coupled receptor (GPCR)-like protein ^19–21^. When bound to Hedgehog PTCH1 is inactivated, permitting SMO to become active and trigger downstream signaling events ^21–24^. Mechanistically, the activation of SMO requires binding of a sterol, likely entering the 7TM bundle from the inner leaflet of the plasma membrane ^25–30^. PTCH1 is proposed to prevent SMO activation by transporting sterols from the inner leaflet of the plasma membrane, thereby limiting SMO access to activating sterols ^31–34^. A hydrophobic conduit coursing through the PTCH1 extracellular domain is required for this transport activity ^33^, and Hedgehog blocks this conduit and inactivates PTCH1 by inserting its essential amino-terminal palmitoyl adduct ^32–35^.

Transporters typically act by moving through a repeated cycle of conformational changes. If PTCH1 transport function employs such a conformational cycle, an agent that preferentially binds and stabilizes a specific PTCH1 conformation would be expected to disrupt its conformational cycle and transport activity, thus permitting activation of SMO. Such an agent thus may serve as a pathway modulator that could make lipid modifications dispensable and may shed light on conformational changes that occur during the PTCH1 working cycle.

## Development of a conformation-specific pathway activating agent

Nanobodies are single-chain antibody fragments ^36^ that have frequently been used to stabilize specific GPCR protein conformations ^37^, and are amenable to genetic engineering. We have chosen as a starting point a synthetic yeast display library to select for conformation-specific nanobodies against PTCH1 ^38^. To select conformation-specific nanobodies we first introduced conformational bias in PTCH1 by altering three acidic residues buried within its transmembrane domain (termed PTCH1-NNQ). These acidic residues, conserved within the RND transporter family, are required for PTCH1 activity in sterol transport and SMO regulation ^26, 33^ and are more generally proposed to drive conformational changes in RND transporters in response to cation influx (Fig. 1a). We thus reasoned that alteration of these residues in PTCH1 might affect the relative representation of its conformational states.

**Figure 1.**
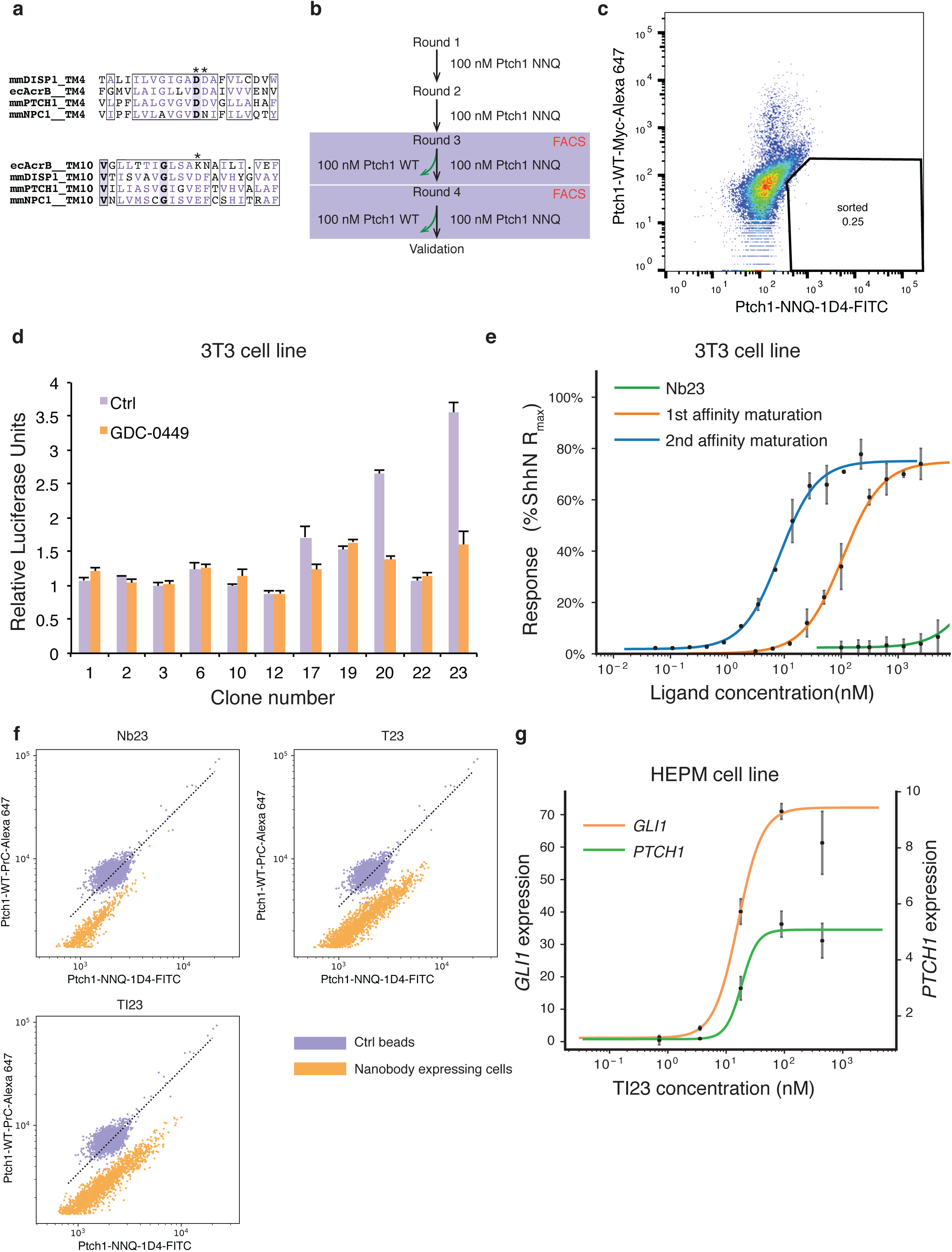
Selection of conformation-selective nanobodies. (a) Alignment of transmembrane 4 and 10 from different RND transporters. The charged residues are marked by asterisks. (b) Flow chart of the steps for nanobody selection. The yeast library was first enriched with MACS for clones that bind to PTCH1-NNQ variant and then the population that prefers the NNQ variant was selected in FACS using PTCH1-NNQ and PTCH1-WT with different fluorescent labels. (c) Yeast cells stained with PTCH1-NNQ (FITC label) and PTCH1-WT (Alexa 647 label) are shown in the FACS plot. In the lower right quadrant are the cells that prefer NNQ variant to the WT variant. Due to more non-specific binding to Alexa 647 fluorophore than the FITC fluorophore, the double positive population shifts towards the upper left quadrant. (d) Nanobodies expressed and purified in *E. coli* were tested on Hedgehog-responsive 3T3 cells with a Gli-dependent luciferase reporter. GDC-0449, a pathway antagonist, is a control showing that nanobodies 17, 20 and 23 display weak activation in this assay. (e) Initial nanobody sequences of clones 17, 20 and 23 were mutagenized and selected in yeast display to obtain higher affinity clones (affinity maturation). After two rounds of affinity maturation, the new nanobody variant, named TI23, exhibits an EC50 of 8.6 nM in 3T3 cells, close to that of the native Hedgehog ligand. (f) The TI23 clone resulting from two rounds of affinity maturation showed a preference for binding to PTCH1-NNQ variant. Yeast cells expressing Nb23, T23 or TI23 were incubated with a mixture of 1:1 Protein C tagged PTCH1-WT and 1D4 tagged PTCH1-NNQ proteins, and then stained with antibodies against protein C tag or 1D4 tag. OneComp beads were used as a control for non-selective binding, as these beads bind to the constant region of kappa chain, and do not discriminate between different antibodies used for staining. (g) In the human mesenchymal cell line HEPM, TI23 activated Hedgehog response and induced transcription of pathway targets, *GLI1* (EC_50_ = 16.0 nM) and *PTCH1* (EC_50_ = 18.5 nM), as assayed by qPCR.

We used purified PTCH1-NNQ variant protein for selection of nanobody clones from the yeast display library. After several rounds of enrichment for PTCH1-NNQ binding yeast clones, we selected nanobodies that preferentially bind to PTCH1-NNQ versus wild-type PTCH1, using FACS (Fluorescence Activated Cell Sorting) and wild-type and NNQ PTCH1 proteins labeled with antibodies coupled to different fluorophores (Fig. 1b). Yeast cells expressing preferentially-bound nanobodies form a population off the diagonal of the FACS plot (Fig. 1c). After selecting nanobody-expressing yeast cells in the NNQ-preferring population, 15 unique clones were identified from sequencing, of which three were discarded because they bind directly to the antibody used during selection (Extended Data Fig. 1a, b). As PTCH1 and PTCH1-NNQ differ only in the acidic residues in the transmembrane domain, differences in nanobody binding most likely derive from differences in conformational states between PTCH1 and PTCH1-NNQ.

Stabilization of a specific PTCH1 conformation would be expected to inactivate its transport activity and permit downstream response in the Hedgehog pathway. We therefore tested the activity of purified nanobody proteins on 3T3-Light2 cells, using a Gli-dependent luciferase assay. Clones 17, 20, and 23 showed weak activation effects (Fig. 1d). We enhanced signaling potency through two rounds of affinity maturation, first by selection from an error-prone PCR library (Extended Data Fig. 1c, d), and then from a library targeting the complementarity-determining regions (CDRs) using one-pot mutagenesis ^39^(Extended Data Fig. 1e, f). The first round of affinity maturation yielded a series of nanobody clones deriving from clone 23, with H105R, G106R substitutions in CDR3 and several variant residues at G50 in CDR2. Only one of these variants, the G50T substitution (named “T23”), could be expressed for purification from *E. coli.* T23 showed enhanced potency in Gli-dependent luciferase assays (Fig. 1e), and was used as the starting sequence for a second round of affinity maturation, in which all CDR residues were systematically randomized in one-pot mutagenesis. After selection based on PTCH1 binding, Y102I in CDR3 was enriched, as well as T77N, an unintended substitution. The purified TI23 nanobody also exhibited greater potency in pathway activation than its T23 parent (Fig. 1e). All of the nanobody variants showed preferential binding for PTCH1-NNQ, as revealed by two-color staining of yeast cells expressing these variants (Fig. 1f; Extended Data Fig. 1g). TI23 also strongly activated human Hedgehog pathway targets *GLI1* and *PTCH1* at low nanomolar concentrations when tested in a cell line derived from human embryonic palatal mesenchymal (HEPM) ^40^(Fig. 1g).

## *In vivo* activation of the Hedgehog pathway

As a small protein (∼12kDa), a nanobody might be expected to display excellent tissue penetrance and be readily accessible to cells in most tissues. We tested the activity of TI23 by intravenously injecting mice with adeno-associated virus (AAV) engineered to express it. This experiment should permit observation of biological effects elicited by sustained nanobody exposure as AAV infection is maintained over several weeks. We monitored lingual epithelium and skin, as these tissues display well-characterized responses to Hedgehog pathway activation ^10, 41^.

We examined *Gli1* mRNA by fluorescence *in situ* hybridization (FISH) as an indicator of pathway activation in lingual epithelium, and noted marked expansion of the range of *Gli1* expression in ShhN or TI23-virus injected mice as compared to the animals that received the control virus (Fig. 2a). A similar expansion of *Gli1* expression was also noted in mice given SAG21k (Fig. 2a), a small molecule Hedgehog agonist that activates SMO ^13^.

**Figure 2.**
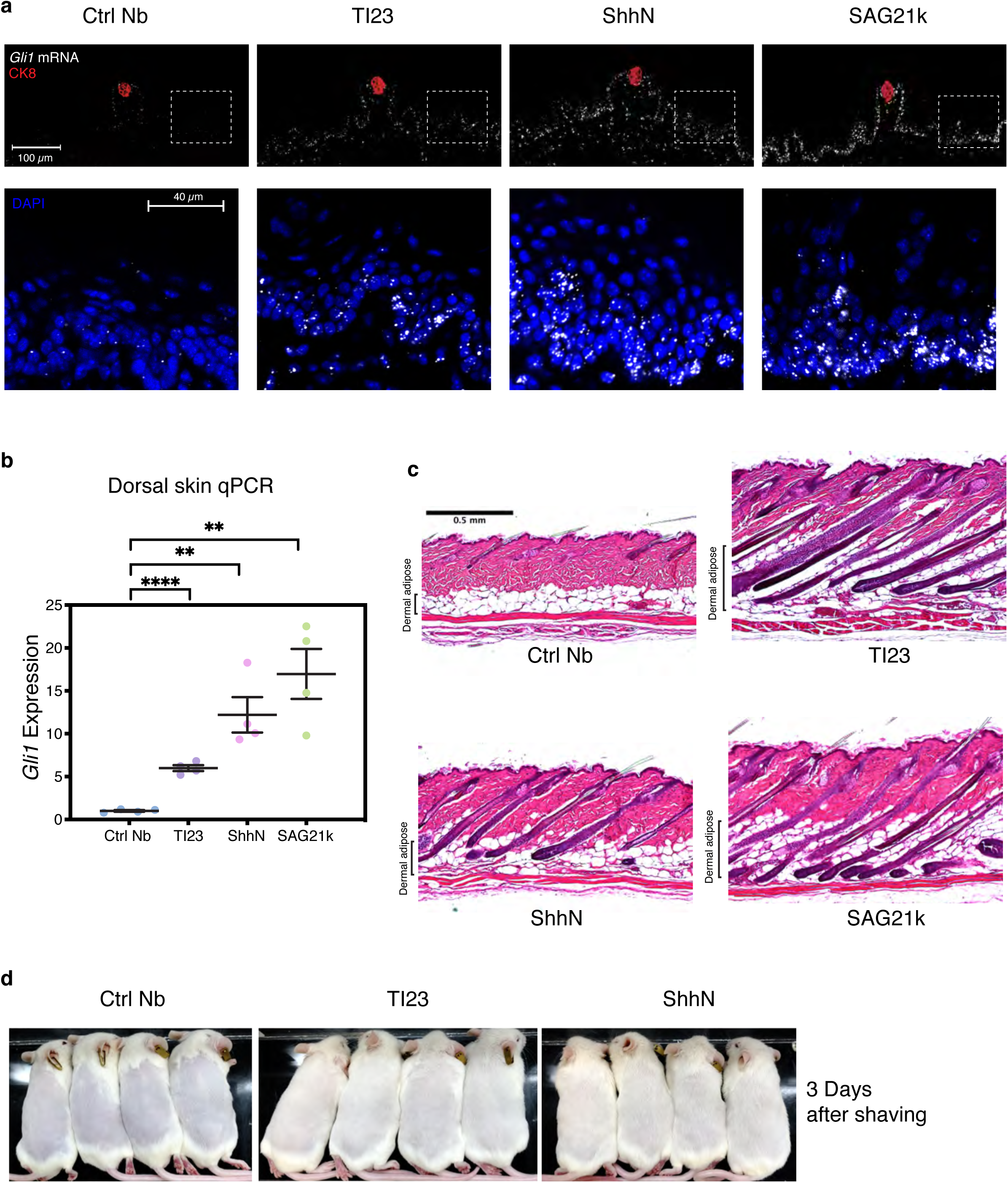
Validation of TI23 activity *in vivo*. (a) TI23 induced *Gli1* expression in lingual epithelial cells, as indicated by *in situ* hybridization using the RNAscope method. Animals receiving AAV encoding different nanobody sequences or ShhN were sacrificed 2 weeks after injection. *Gli1* expression is limited to epithelial cells immediately surrounding the taste receptor cells marked by expression of CK8, whereas pathway agonists, TI23, ShhN, or the small molecule SAG21k, expanded the range of response to almost the entire epithelium, as shown in the inset panels. (b) *Gli1* expression was activated in the dorsal skin of animals receiving TI23, ShhN or the small molecule SAG21k, suggesting that TI23 activated the Hedgehog pathway in the skin. (c) Histology of the dorsal skin suggests that hair follicles in the control group are in quiescent telogen phase, whereas hair follicles grow and invade the adipocyte layer in with TI23, ShhN, or SAG21k treatment, indicating induction of anagen. (d) Hair regrowth observed two weeks after virus injection is much accelerated in TI23 or ShhN-treated animals as compared to the control group, suggesting that these hair follicles are in active anagen phase. N=4 in each group used in (a-c), and n= 4 in each group used in (d).

The TI23 nanobody also augmented Hedgehog pathway activity in the dorsal skin, as indicated by a 6-fold increase in *Gli1* RNA levels (Fig. 2B). We also noted lengthening hair in the hair follicle and expansion of dermal adipocytes upon histological examination of dorsal skin in mice infected with AAV encoding TI23 or ShhN, but not control nanobody, indicating hair follicle entry into the anagen phase of the hair cycle (Fig. 2c). Consistent with accelerated entry into anagen, we noted faster hair regrowth on the dorsal skin after shaving (Fig. 2d).

## Structure of the PTCH1::TI23 complex

To determine the conformational effects of TI23 binding to PTCH1 we prepared the PTCH1::TI23 complex for structure determination by cryo-EM. The complex was clearly visualized in cryo-EM micrographs (Extended Data Fig. 2a), with well-fitted contrast transfer function parameters (CTF; Extended Data Fig. 2b) and 2D class averages (Extended Data Fig. 2c). 3D reconstruction of a cryo-EM dataset yielded a high quality map (Fig. 3a; procedure in Extended Data Fig. 2d) at a resolution of 3.4 Å (Extended Data Fig. 2f; Table 1). All 12 transmembrane (TM) helices and two major extracellular domains (ECDs) were resolved (Fig. 3a), and an atomic model of the PTCH1::TI23 complex was built based on this map and the previously determined murine PTCH1 structure ^33^. Most of the intracellular sequence was unresolved, and not modeled, except for two transverse helices preceding TM1 and TM7 (Fig. 3b).

**Figure 3.**
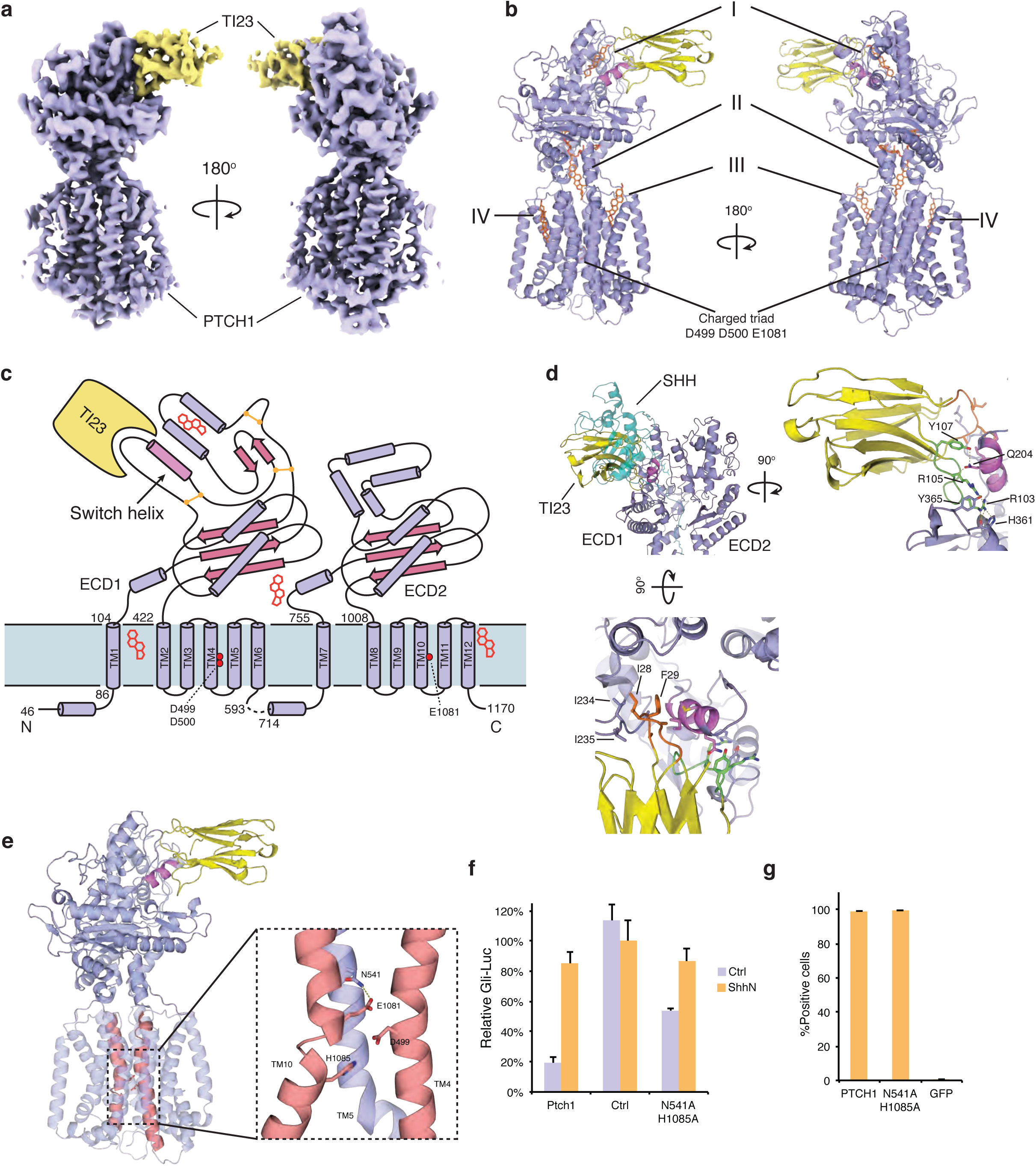
Overview of mouse PTCH1::TI23 complex structure. (a) The cryo-EM map of PTCH1::TI23 complex shows clear features of the proteins. PTCH1, violet; TI23, yellow. (b) Protein model of the complex with PTCH1 and TI23 colored as in A. Lipid-like densities found in the map were modeled in sites I through IV. (c) Schematic view of PTCH1 showing the secondary structure elements and the relative positions of TI23 and the lipid-like densities. The key helix involved in the conformational change is highlighted as “switch helix”. (d) The binding site of TI23 on PTCH1 overlaps with that of SHH (teal). The switch helix, highlighted in violet, is sandwiched by CDR1 and CDR3 of TI23. The hydrophobic interactions from CDR1 are viewed from above the membrane, whereas the hydrogen bond interactions from CDR3 are viewed from the ECD2 side of PTCH1 protein. (e) The charged triad is located in TM4 and TM10, the two helices highlighted in red. Binding of TI23 to wild-type PTCH1 improved the resolution of side chains in the transmembrane, suggesting potential allostery between the two parts of PTCH1. Interactions of the charged triad with adjacent residues are shown in the inset. E1081 is stabilized by hydrogen bonding with N541, whereas D499 forms a salt bridge with H1085. (f) Alanine substitution of N541 and H1085 impaired PTCH1 activity, as indicated by Gli-dependent luciferase assays in *Ptch1*^-/-^ cells. (g) PTCH1 N541A H1085A variant still binds to Shh when expressed in 293 cells, suggesting that it preserves the normal fold.

**Table 1.**
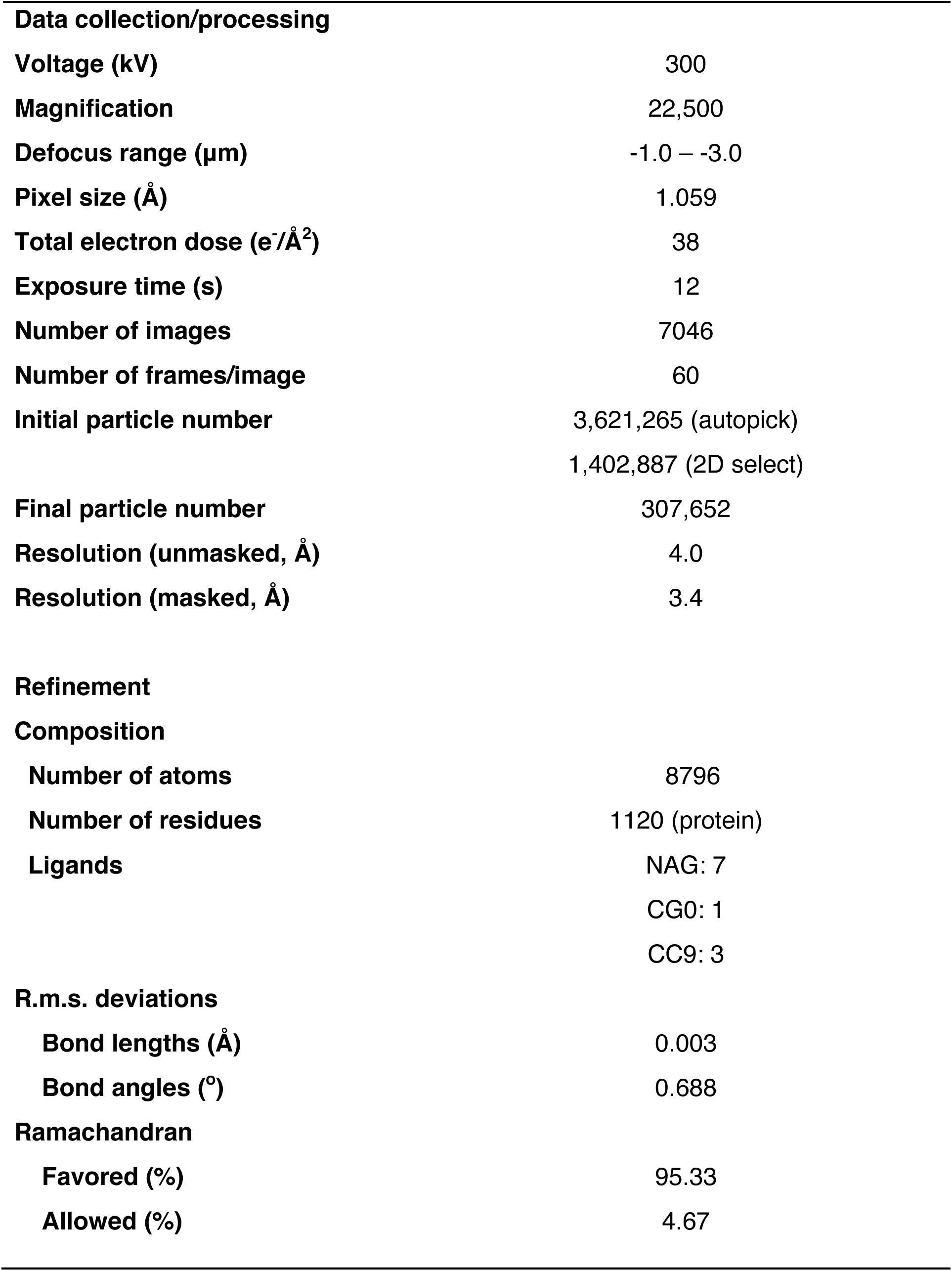
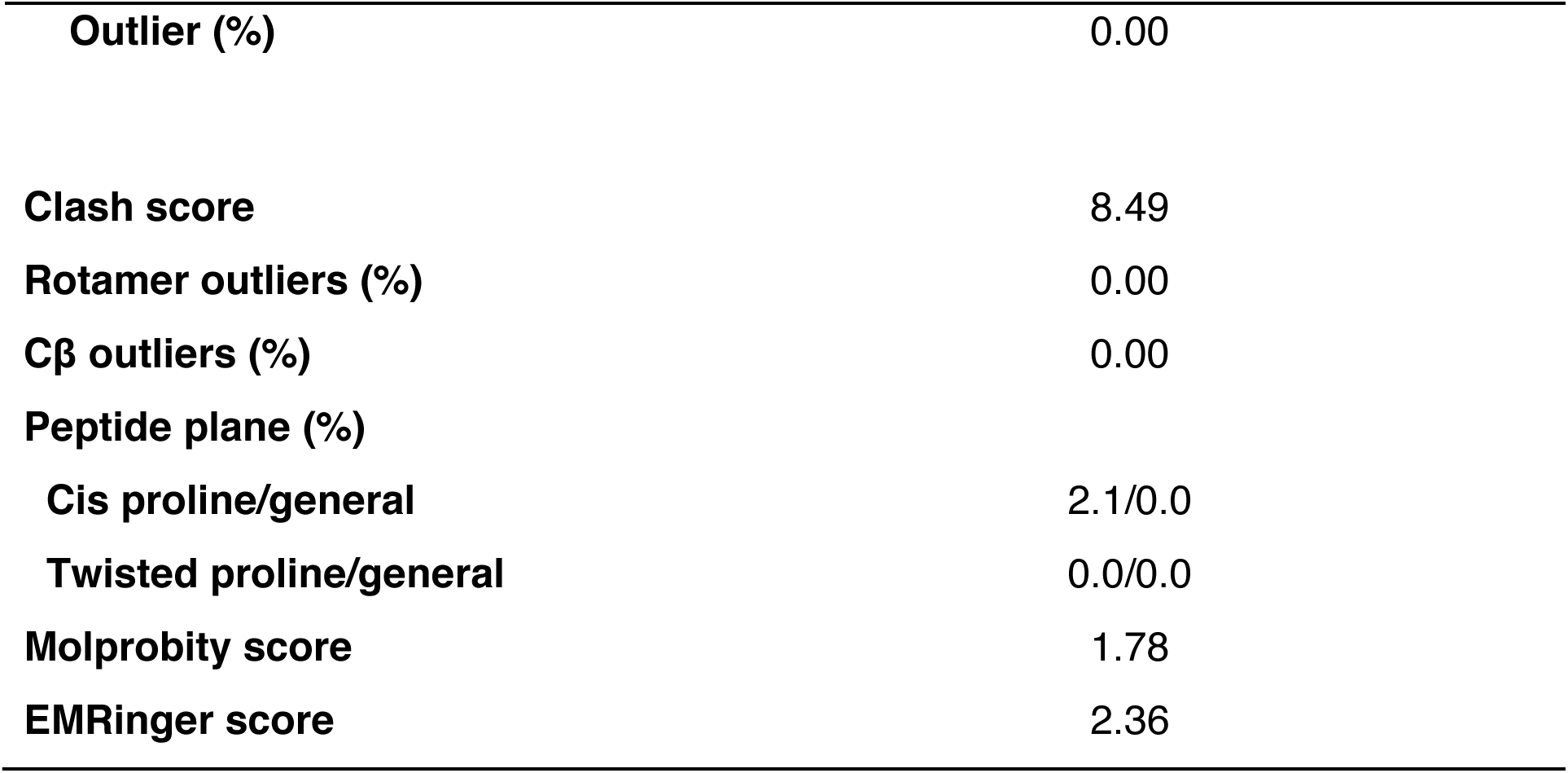
Summary of cryo-EM data collection and model refinement

|  |  |
| --- | --- |
| **Data collection/processing** |  |
| **Voltage (kV)** | 300 |
| **Magnification** | 22,500 |
| **Defocus range (μm)** | -1.0 – -3.0 |
| **Pixel size (Å)** | 1.059 |
| **Total electron dose (e<sup>-</sup>/Å<sup>2</sup>)** | 38 |
| **Exposure time (s)** | 12 |
| **Number of images** | 7046 |
| **Number of frames/image** | 60 |
| **Initial particle number** | 3,621,265 (autopick)<br>1,402,887 (2D select) |
| **Final particle number** | 307,652 |
| **Resolution (unmasked, Å)** | 4.0 |
| **Resolution (masked, Å)** | 3.4 |
| **Refinement** |  |
| **Composition** |  |
| **Number of atoms** | 8796 |
| **Number of residues** | 1120 (protein) |
| **Ligands** | NAG: 7<br>CG0: 1<br>CC9: 3 |
| **R.m.s. deviations** |  |
| **Bond lengths (Å)** | 0.003 |
| **Bond angles (°)** | 0.688 |
| **Ramachandran** |  |
| **Favored (%)** | 95.33 |
| **Allowed (%)** | 4.67 |
| **Outlier (%)** | 0.00 |
| **Clash score** | 8.49 |
| **Rotamer outliers (%)** | 0.00 |
| **Cβ outliers (%)** | 0.00 |
| **Peptide plane (%)** |  |
| **Cis proline/general** | 2.1/0.0 |
| **Twisted proline/general** | 0.0/0.0 |
| **Molprobity score** | 1.78 |
| **EMRinger score** | 2.36 |

Sterol-like densities were identified in multiple sites, one in a pocket at the distal tip of ECD1 (farthest from the membrane, density I), one in the cavity proposed as part of the transport conduit (II) and two more at the periphery of the transmembrane domain (III and IV) (Fig. 3b). The density in site II is especially well resolved, and its unusual “Y” shape strongly suggests that it is glyco-diosgenin (GDN), the detergent used during sample preparation (Extended Data Fig. 3b). The other sterol-like densities are also most likely GDN, but only the steroidal moiety of GDN, digitogenin, was resolved and modeled.

The nanobody interacts only with ECD1 of PTCH1, as shown in the schematic drawing (Fig. 3c). The binding site of TI23 overlaps with that of SHH, which interacts with both ECD1 and ECD2. CDR1 and CDR3 loops from TI23 sandwich a short helix in PTCH1 ECD1, which we name as the “switch helix”. CDR1 primarily interacts with PTCH1 by inserting hydrophobic residues I28 and F29 into the hydrophobic pocket at lipid site I, whereas CDR3 primarily forms a hydrogen bond network with other residues on the surface of PTCH1 (Fig. 3d).

Although TI23 interacts exclusively with ECD1, we noted significant improvement in the resolution of side chains within the transmembrane domain. Of particular interest, the charged residue triad within the transmembrane (TM) domain that was altered for selection of TI23 is clearly visible as compared to other published PTCH1 structures, none of which have well resolved side chain density for these residues. This apparent stabilization of transmembrane residues suggests potential communication of the transmembrane domain with the ECD. Such allostery would account for the improvement of the TM resolution upon TI23 binding to wild-type PTCH1, while also explaining preferential binding of TI23 to PTCH1-NNQ during selection (Fig. 1c). Our model further suggests that these acidic residues may be stabilized by side chain interaction with residues nearby, namely, N541 in TM5 and H1085 in TM10 (Fig. 3e). Indeed, alanine substitution of these residues impaired PTCH1 activity (Fig. 3f) without affecting the overall fold, as indicated by preserved binding to ShhN (Fig. 3g).

The overall structure of the PTCH1::TI23 complex is similar to the unbound murine PTCH1 structure, with a root mean square deviation of 0.955 Å of the Cα carbon atoms over 910 residues. Both ECD1 and ECD2 display some conformational differences in the complex. One minor difference is a rotation of ECD2 around its connection to the TM domain by ∼5 degrees towards ECD1 as compared to PTCH1 alone (Fig. 4a). A more marked difference is the rotation by ∼32 degrees of the distal tip of a short helix within ECD1 towards the membrane, in a manner suggestive of a flipped switch (Fig. 4a, inset). We term this helix the “switch helix” (residues 203-209, highlighted in Fig. 3c), and refer to its conformations in PTCH1 alone and in the PTCH1::TI23 complex as poses 1 and 2, respectively. These two alternative poses of the switch helix are present but have gone unremarked in other structures of PTCH1 determined under various conditions. For example, in the ternary complex of a single native Shh ligand bound to two human PTCH1 molecules ^32^, PTCH1 from chain a, the molecule whose sterol conduit is occluded by interaction with the N-terminal palmitoyl moiety of the SHH ligand, adopts pose 2, whereas PTCH1 from chain b adopts pose 1 ^32^. Indeed, in all published structures of PTCH1 the switch helix adopts one or the other of these two poses^15, 31–35, 42^, suggesting that they represent discrete alternative conformations preferentially populated within the PTCH1 activity cycle (Fig. 4b).

**Figure 4.**
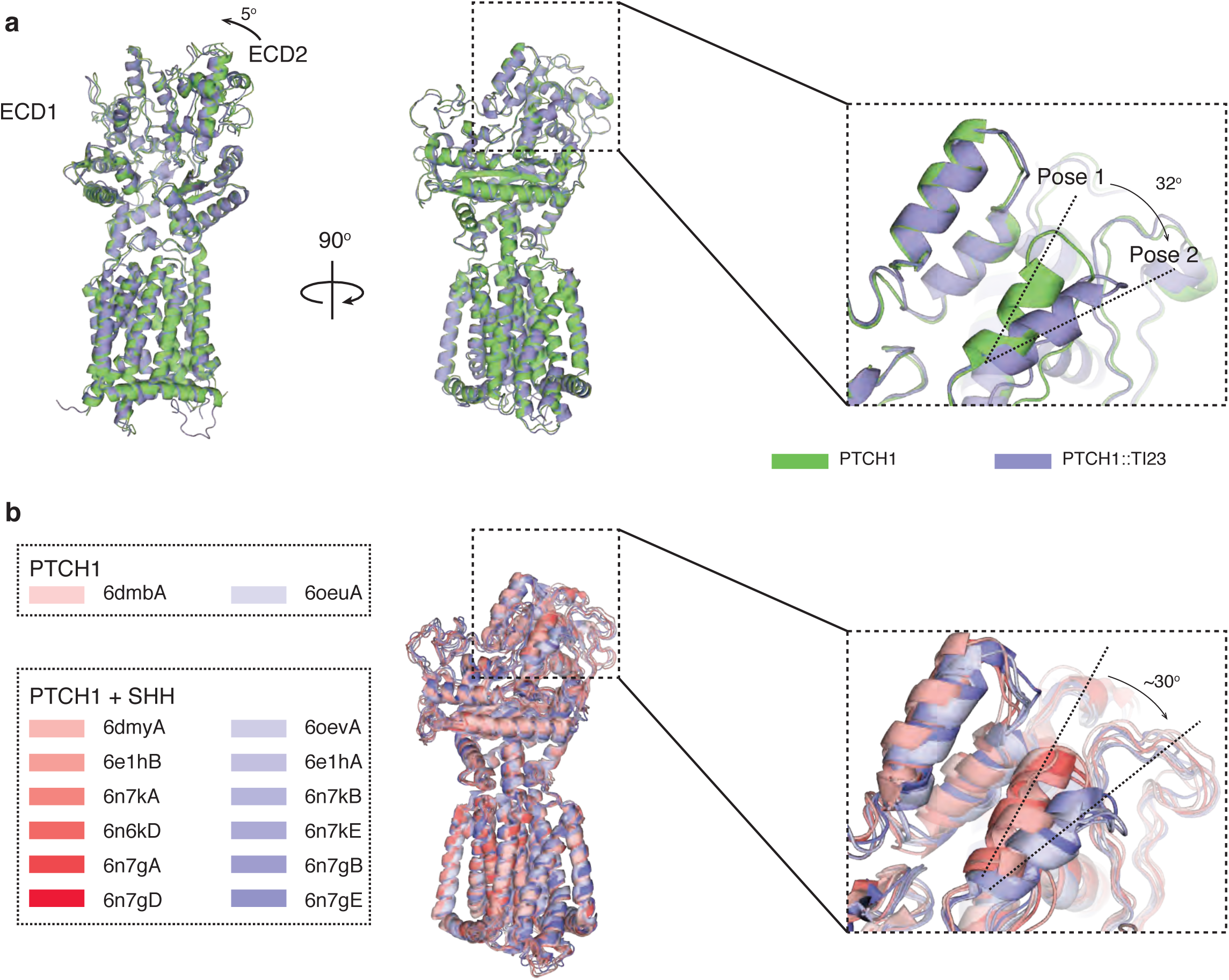
Conformational change induced by TI23. (a) Overlay of the structures of murine PTCH1 alone or in complex with TI23 shows two major changes in the extracellular domain. The extracellular domain 2 between TM7 and TM8 turns around 5° pivoting on its connection to the transmembrane domain. A short helix (the switch helix) in extracellular domain 1 rotates ∼32° towards the membrane. The conformation in PTCH1 alone and in the complex is referred to as pose 1 and 2, respectively. (b) Other published PTCH1 structures also fall into Pose 1 and 2 categories. In this overlay of all the structures, pose 1-like structures are shown in shades of red, and pose 2-like structures in shades of blue.

## Effects of the switch helix on the sterol conduit

These structural rearrangements alter the shape of the transport conduit as assessed by the Caver program (Fig. 5a). The region of the conduit encompassing sterol I in murine PTCH1 thus is seen to be dramatically constricted in the conduit of the PTCH1::TI23 complex, and the conduit in the PTCH1::TI23 complex also acquires a distal opening to the exterior (Fig. 5b). In parallel with this change in conduit shape, the bound sterol-like density shifts from a more proximal enclosed cavity to a more distal position with an opening to the exterior (Fig. 5c). This concerted proximal constriction and distal expansion results primarily from rotation of the switch helix. If PTCH1 activity is, like other RND family members, driven by a chemiosmotic gradient^26^, the conformational change identified here may form part of a defined sequence that results in directional movement of substrates within the transport conduit.

**Figure 5.**
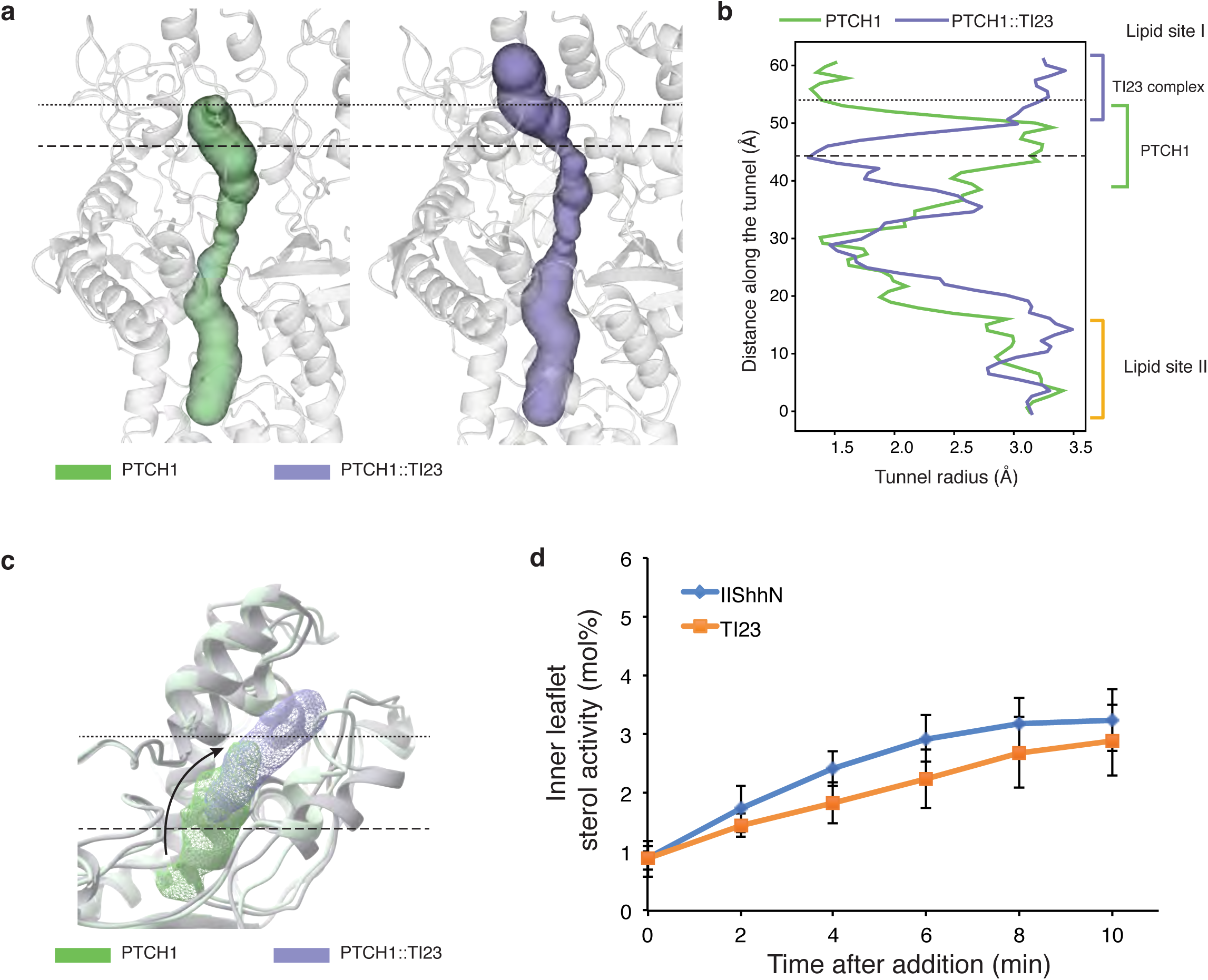
Effect of nanobody-induced conformational changes on the sterol conduit. (a) The rotation of the switch helix alters the shape of the cavity within the extracellular domain. In the PTCH1 structure the conduit is capped at the end, as indicated by the dotted line (…), whereas in the TI23 bound structure, the end of the conduit is wide open to the exterior and the lower part is throttled, as marked by the dashed line (---). (b) The radii at different points along the conduit are plotted here, with the altered parts marked with two vertical lines. TI23 binding opens the upper end of the conduit but closes the lower part of lipid site I. (c) Position of the lipid-like density in site I changes with TI23 binding. The rotation of the switch helix may push the bound substrate outwards while closing down the entry route. (d) In *Ptch1*^-/-^ MEFs transfected with PTCH1, plasma membrane inner leaflet (IPM) cholesterol activity increased immediately after adding purified TI23, or Hedgehog ligand (ShhN). Hedgehog ligand caused a slightly faster increase in IPM cholesterol activity, which plateaued after ∼6 min. This may reflect the difference in efficacy of these two ligands, as TI23 induces ∼75% maximum pathway activity at saturating concentration in Gli-dependent luciferase assays.

The best-studied bacterial RND homolog, AcrB, also undergoes a series of conformational transitions that affect the substrate conduit, similar in principle although distinct in detail from that of PTCH1. As the AcrB transporter moves between conformations, the substrate conduit changes shape to admit substrate proximally and move it to the central portion of the conduit (L and T states), then to open distally while constricting proximally (O state), thus expelling substrate from the distal tip of the ECD ^43^.

The TI23 nanobody appears to stabilize pose 2 of the PTCH1 switch helix. If PTCH1-mediated transport of sterols away from the inner leaflet indeed depends on the dynamic changes in the shape of the conduit associated with switch helix movement, TI23 binding may lock PTCH1 in a state that is incompatible with sterol movement. To test this idea, we utilized a solvatochromic fluorescent sterol sensor, microinjected into cells to permit ratiometric measurement of sterol available for binding within the inner leaflet of the membrane ^44^. This sensor previously revealed that available sterol decreases sharply with PTCH1 activity, and that PTCH1 inactivation by Shh ligand causes a return to normal sterol availability ^33^. Similar to the effect of Shh ligand addition, we noted that TI23 addition reversed the PTCH1-mediated reduction in cholesterol activity (Fig. 5d).

## Discussion

The therapeutic applications of Hedgehog pathway modulation have focused primarily on pathway antagonists, and inhibition of the Hedgehog pathway has proven efficacious in the treatment of cancers driven by excessive Hedgehog pathway activity directly in primary cells of the tumor ^1, 2^. In other cancers, however, Hedgehog pathway blockade may accelerate tumor growth and progression. These include pancreatic, colon, and bladder cancers, and likely other malignancies originating from organs of endodermal origin, in which Hedgehog pathway activity in stromal cells may restrain cancer progression. Other therapeutic applications for pathway activation may include regeneration of taste receptor cells in chemotherapy-treated patients ^10^ or regeneration of bone in diabetic patients ^12^.

Current candidates as pharmaceutical agents that activate the Hedgehog pathway are all hydrophobic in nature, including small molecule members of the SAG family ^45^, certain oxysterols ^46, 47^, and purmorphamine ^48^, all of which target SMO, and the lipid-modified Hedgehog protein or its derivatives, which target PTCH1. Our conformation-selective PTCH1-directed nanobody TI23 represents a new class of potent, more hydrophilic agonists, which unlike the native Hedgehog protein does not require hydrophobic modification for activity. TI23 furthermore has the potential to be engineered for targeting by fusion to an antibody or other agent with tissue or cell-type specificity. This potential for greater specificity is critical for translational applications, as Hedgehog responsive cells are widespread and systemic pathway activation may have pleiotropic effects and cause complications such as fibrosis.

TI23 is not only a promising candidate for further pharmaceutical development, but also provides insight into the PTCH1 transport mechanism. Directional movement of substrate through a transporter protein implies conformational change, but the identification of such conformational transitions for transporters is a nontrivial challenge. Our conformation-specific nanobody approach allowed us to identify two distinct conformations associated with poses 1 and 2 of the PTCH1 switch helix. The changes in shape of the transport conduit associated with these poses suggest a potential mechanism for directed substrate movement. As PTCH1 is distinct from the well-characterized RND transporter AcrB in both its preferred substrate and its extracellular domain structure^49^, it is not surprising that the conformational transitions of these proteins differ. Indeed, given these differences, the apparent similarity in peristaltic movement of the substrate conduit in both proteins seems quite remarkable.

Our conformation-selective nanobody approach may be generalizable to the study of other transporters, in particular other members of the RND family. In mammals this family includes the NPC1 cholesterol transport protein ^50^, and other PTCH-like proteins, such as PTCHD1, disruption of which is strongly associated with autism ^51, 52^. For other transporters, mutations that disrupt function may do so by biasing the normal conformational landscape without uniquely stabilizing any one conformation. Selection of nanobodies that preferentially bind such mutants may enable capture of sparsely populated yet critical conformations, expanding the repertoire of experimentally accessible states for structural and functional studies.

## Materials and Methods

### Cell culture

Sf9 and 293T cells were maintained in culture according to previously published conditions (Myers et al., 2013). 293-Freestyle cells were maintained in suspension culture in an 8% CO_2_ incubator equipped with a shaking platform, using Freestyle 293 expression medium (Life Technologies) supplemented with 1% fetal bovine serum (Gemini Bio). Baculovirus production in Sf9 cells and infection of suspension 293 cultures with recombinant baculovirus (BacMam expression) was performed as previously described (Myers et al., 2013). 3T3 cells and *Ptch1*^-/-^ MEFs were maintained as previously described (Myers et al., 2017).

### Molecular cloning

All constructs were cloned with Gibson assembly. For BacMam expression, PTCH1 variants were cloned into pVLAD6 vector. For yeast selection, Ptch1-C and Ptch1-C-NNQ variants were used. Ptch1-C is mouse PTCH1 truncated at amino acid 1173, deleted at 619-711 and altered at C1167Y. Use of Ptch1-C for selection minimized the possibility of getting nanobodies that bind to PTCH1 intracellular domain, due to extensive deletion of the intracellular sequence. For structural determination and cell biology experiments, Ptch1-B as reported earlier was used. For luciferase assay and cell surface binding experiments, PTCH1 variants were cloned into pcDNA-h (pcDNA3 vector with the neomycin resistance cassette removed).

### Yeast display selection

The synthetic nanobody library was grown in SDCAA media at 30 C to a cell density of ∼1×10^8^/ml. Cells covering about 10 times the initial diversity (5×10^8^ diversity, 5×10^9^ cells) were transferred into SGCAA media at 20C to induce expression of nanobody on cell surface. For selection, 7.5×10^9^ cells were pelleted by centrifugation and resuspended in selection buffer (20 mM HEPES, pH 7.5, 150 mM NaCl, 0.5 mg/ml BSA, 0.1% DDM, 0.02% CHS). The cells were then incubated with 100 nM 1D4-tagged Ptch1-C NNQ, spun down and washed with selection buffer, and then with FITC-labeled 1D4 antibody, then 100 µL anti-FITC MACS beads. After loading the beads-bound cells onto the magnetic manifold and washed extensively with selection buffer, the bound cells were eluted, cultured in SDCAA media and induced for nanobody expression in SGCAA media. A second round of selection was then performed on these cells, first with the Alexa647 labeled 1D4 antibody alone to counter-select antibody-binding cells and then with 100 nM 1D4 tagged Ptch1-C NNQ. The selected cells were grown in SDCAA and incuded with SGCAA again and then incubated with 100 nM Myc-tagged Ptch1-C and 100 nM 1D4-tagged Ptch1-C-NNQ and stained with anti-Myc Alexa 647 and anti-1D4 FITC and cells showing stronger FITC signal on FACS were selected. The same FACS selection was repeated and the selected cells were grown and dilution-plated. Plasmid was prepared from single colonies and sequenced after rolling cycle amplification (RCA). 15 unique sequences were retrieved from 24 colonies. Yeast cells harboring these nanobody sequences were then tested for binding to anti-1D4 antibody and to Ptch1-C-NNQ. Three out 15 bind to 1D4 antibody directly and were excluded from later analysis. The rest of the sequences were cloned into pET28a vectors for expression and purification from *E. coli*.

### Nanobody purification

pET26b vectors containing nanobody sequences were transformed into *E. coli* BL21 (DE3) strain. The bacteria were grown in Terrific broth media at 37 °C to OD600 of 0.8, and then induced with 0.2 mM IPTG and transferred to 20 °C. After overnight expression, the cells were harvested by centrifugation at 8,000 g. The cell pellet was resuspended in SET buffer (500 mM sucrose, 0.5 mM EDTA, pH 8.0, 200 mM Tris, pH 8.0) at a ratio of 5 ml buffer /1 g pellet. After stirring for 30 min at room temperature, two volumes of water was added. After stirring for an addition 45 min, MgCl_2_ was added to 2 mM and benzonase at 1:100,000. After 5 min incubation, NaCl was added to 150 mM, imidazole to 20 mM and the whole mixture was centrifuged at 20,000 g for 15 min at 4 C. The supernatant was then loaded onto a Ni-NTA column, washed with ice-cold buffer (20 mM HEPES pH 7.5, 500 mM NaCl, 20 mM imidazole) and then eluted in 20 mM HEPES pH 7.5, 150 mM NaCl, 250 mM imidazole. The eluted protein was then dialyzed overnight in 20 mM HEPES pH 7.5, 150 mM NaCl at 4 °C. All of the initial hits except for clone 13 could be expressed and purified. Clone 13 was then excluded from analysis.

### Affinity maturation

The first round affinity maturation library was made with error-prone PCR. Nanobody clone 17, 20 and 23 were chosen as the starting point of this selection. 10 ng plasmid containing the nanobody sequence was used as the template (equivalent to ∼1 ng DNA of nanobody sequence) and PCR amplified with Mutazyme kit. The PCR product was gel-purified and 10ng was then used as the template for the next round of PCR. A total of 4 rounds of PCR were performed. The final product was then amplified with Phusion polymerase to obtain sufficient amounts for yeast transformation. A total of ∼100 µg DNA was purified for each parental sequence using ∼2 µg of the error-prone PCR product. The DNA fragments were then transformed into yeast along with pYDS2.0 plasmid backbone. DNA from 3 different parental sequence, and a mixture of the three were electroporated separately into yeast cells, but the cells were pooled in YPD for recovery after electroporation. Serial dilution and plating gave an estimate of 1×10^9^ independent transformant for this library. The transformed yeast cells were then grown in YPD media with 100 µg/ml nourseothricin sulfate, and then induced in YPG media with the same antibiotic. The yeast cells were enriched for PTCH1 binding by MACS selection using concentrations of 1D4-tagged Ptch1-C NNQ at 100 nM, 5 nM, 0.8 nM. Then cells expressing nanobody were incubated with Ptch1-C NNQ at 0.6 nM. After washing in selection buffer, the cells were incubated with the parental 17, 20, 23 nanobody proteins at 1 µM each for 170 min at room temperature. The cells were then stained with FITC-labeled HA antibody to mark nanobody expression levels and Alexa 647-labeled anti-1D4 antibody to mark PTCH1 binding. Cells that maintain high PTCH1 binding were selected from FACS. 64 clones were sequenced to identify repeating changes.

The second round of affinity maturation was performed with a library targeting the complementarity determining regions (CDRs) using the one-pot mutagenesis method (Wrenbeck et al., 2016). A pool of DNA oligos with NNK substituting each codon in the CDR regions was used for one-pot mutagenesis of the CDRs so that theoretically all 20 amino acids at each position were represented in this library. The DNA product from one-pot mutagenesis was then amplified with Q5 polymerase and purified with gel extraction. A final product ∼5 µg DNA was used for yeast transformation. The transformed cells were grown in YPD media containing 100 µg/ml nourseothricin sulfate and induced in YPG media containing the same antibiotic. The cells were then incubated with 10 nM protein C-tagged Ptch1-C, washed in selection buffer and then incubated with 1µM 23T (purified nanobody protein with the consensus sequence from the 1^st^ round of affinity maturation) for one day. The cells were then stained with FITC-labeled HA and Alexa 647 labeled anti-protein C antibody and the PTCH1-high cells were selected in FACS. The cells were grown in YPD and induced again. The same FACS selection procedure was repeated to further purify the population. The nanobody sequences from the plasmids prepared from the initial yeast library and the final selected library were then amplified with Q5 polymerase and sent for amplicon sequencing at MGH sequencing core.

### PTCH1 purification

Purification of PTCH1 was performed as previously described with minor changes. Suspension 293 cells were grown to a density of 1.2 – 1.6 × 10^6^/ml, supplemented with 10 mM sodium butyrate, and infected with high-titer Ptch1-SBP baculoviruses for 40-48 hr. Cell pellets were stored at −80°C. Pellets were thawed into hypotonic buffer (20 mM HEPES pH 7.5, 10 mM MgCl_2_, 10 mM KCl, 0.25 M sucrose) supplemented with protease inhibitors and benzonase. Crude membranes were pelleted with centrifugation (100,000 x g, 30 min., 4°C). The pellet was resuspended in lysis buffer (300 mM NaCl, 20 mM HEPES pH 7.5, 2mg/ml iodoacetamide, 1% DDM / 0.2% CHS) with protease inhibitors and solubilized for 1 hour at 4°C with gentle rotation. After centrifugation (100,000 x g, 30 min., 4°C), the supernatant was incubated with streptavidin-agarose affinity resin in batch mode for 2-3 hours at 4°C with gentle rotation. The resin was packed into a disposable column, and washed with 20-30 column volumes of buffer (20 mM HEPES pH 7.5, 300 mM NaCl, 0.03% DDM / 0.006% CHS). Protein was eluted in the same buffer supplemented with 2.5 mM biotin.

### Cryo-EM data acquisition

Eluted Ptch1-B protein was mixed at 1:1.1 ratio with TI23 and then loaded onto Superdex 200 column pre-equilibrated with SEC buffer (20 mM HEPES, pH 8, 150 mM NaCl, 0.02% GDN). The peak fractions were collected and concentrated with an Amicon filter with molecular weight cutoff of 100 kDa to A280 ∼4.5. 2.5 µL sample was applied to a glow-discharged quantifoil grid on a vitrobot. The sample chamber was kept at 100% relative humidity. The grid was blotted for 10s and plunged into liquid ethane bath cooled by liquid nitrogen.

The cryo grids were imaged on a Titan Krios 2 electron microscope operated at 300 kV. Images were taken on the pre-GIF K2 camera in dose fractionation mode, at nominal maginification of 22.5k, corresponding to a pixel size of 1.059 Å (0.5295 Å per super-resolution pixel). The dose rate was ∼8e/pix/sec with a total exposure time was 12s at a frame rate of 0.2s/frame. Fully automated data collection was performed with SerialEM, with a defocus range of −1 µm to −3 µm. Gain reference was taken at the beginning of the data collection and was applied later in data processing.

### Image processing

A total of 7,046 movie stacks were collected. The movie stacks were corrected by gain reference, binned by 2, and corrected for beam-induced motion with MotionCor2. CTF was determined with CTFFIND4 from the motion-corrected sums without dose-weighting using a wrapper provided in cryoSPARC2. Dose-weighted sums were used for all the following steps of processing. Particles were autopicked cryoSPARC2. Particles corresponding to protein molecules were selected from 2D classification. These particles were then reconstructed ab initio, and then classified with heterogeneous refinement into 3 classes, using two copies of the map generated from the last step plus one junk map as the initial models. The best class was chosen for homogeneous refinement and then non-uniform refinement to obtain a map at 4.1 Å. The particles were then analyzed with the 3D variability analysis tool and the two extremes of the first eigenvector were used as the basis for further 3D classification. The final 3D class was refined with non-uniform refinement to a resolution of 3.7 Å. The particle stack was then exported to cisTEM using the scripts in pyEM. After one iteration of local refinement with a mask excluding the detergent micelle, a map was reported at 3.4 Å. The final map after sharpening was used for model building.

### Protein model building

Nanobody TI23 structure was generated with rosettaCM using 4mqtB and 5m30F as the template structures. The generated structure and the previously determined PTCH1 structure (6mg8) were docked into the cryo-EM map and refined in phenix.real_space_refine with morphing. The refined model was then edited manually in coot, to add in residues that are now resolved in the new structure, and the small molecules. The constraints for small molecules were generated on the PRODRG server. The entire structure was then refined in phenix.real_space_refine.

### FACS-based ShhN binding assay

293 cells were transiently transfected with GFP-tagged Ptch1 constructs. After 24 hours, cells were dissociated using 10 mM EDTA, washed with HPBS 0.5 mM Ca^2+^, and pelleted by centrifugation. Cells were then resuspended in binding buffer (HPBS, 0.5 mM Ca^2+^, 0.5 mg/ml BSA) and incubated with purified ShhN-biotin (1:400 dilution) for 30 minutes at 4° C. Cells were then washed three times in binding buffer by centrifugation and subsequently incubated with Alexa Fluor 647 streptavidin conjugate (Invitrogen) for 15 minutes at 4° C. Cells were then washed three times by centrifugation in wash buffer (binding buffer plus 1 mM biotin) and the percentages of cells bound by ShhN were quantified by flow cytometry after gating for PTCH1-GFP expression (BD FACSAria II, Stanford Stem Cell Institute FACS Core).

### Gli-dependent luciferase assay

The luciferase assay was performed in Ptch1^-/-^ MEFs, as previously described(Myers et al., 2017). Ptch1^-/-^ MEFs were seeded into 24-well plates and then transfected with various plasmids along with a mixture containing 8xGli firefly luciferase and SV40-renilla luciferase plasmids. For each well, 2ng (0.4%) plasmid encoding Ptch1-B variants, or 5ng (1%) plasmid encoding full-length PTCH1 was used. When cells were confluent, they were shifted to DMEM with 0.5% serum containing ShhN-conditioned medium or control medium and incubated for 48 hr. Luciferase activity was then measured using a Berthold Centro XS3 luminometer. The ShhN conditioned medium was prepared from 293 cells transfected with a plasmid expressing the amino signaling domain of Shh. In brief, 293 cells were transfected with the ShhN expression plasmid with lipofectamine 2000. Twelve hours after transfection, culture medium was replaced with 2% FBS low-serum medium. The conditioned medium was then collected 48hours after medium change, and used at 1:10 for the luciferase assays.

### Cellular cholesterol measurement

The Perfringolysin O D4 domain (a.a. 391–500) and mutants were expressed as His_6_– tagged proteins in *E. coli* BL21 RIL codon plus (Stratagene) cells and purified using the His_6_–affinity resin (GenScript). These proteins were labeled at the single Cys site (C459) by a solvatochromic fluorophore to generate ratiometric sensors.

Ptch1^-/-^ MEFs were seeded into 50 mm round glass–bottom plates (MatTek) and grown at 37°C in a humidified atmosphere of 95% air and 5% CO_2_ in Dulbecco’s modified Eagle’s medium (DMEM) (Life Technologies) supplemented with 10% (v/v) fatal bovine serum (FBS), 100 U/ml penicillin G, and 100 μg/ml streptomycin sulfate (Life Technologies).

After attachment to the culture vessels (∼24hr), cells were transiently transfected with plasmids encoding Ptch1-B variants using the jetPRIME transfection reagent (Polyplus Transfection) according to the manufacturer’s protocol. 1 µg plasmid was used for each transfection. Cholesterol in the inner (IPM) leaflets of the plasma membrane was quantified using cholesterol sensors as described previously (Liu et al., 2017) with some modification. Specifically, the Y415A/D434W/A463W (YDA) mutant of the D4 domain labeled with (2Z,3*E*)-3-((acryloyloxy)imino)-2-((7-(diethylamino)-9,9-dimethyl-9*H*-fluoren-2-yl)methylene)-2,3-dihydro-1*H*-inden-1-one (WCR) was delivered into the cells by microinjection for quantification of IPM cholesterol ([Chol]*_i_*). All sensor calibration, microscopy measurements, and ratiometric imaging data analysis were performed as described (Liu et al., 2017; Liu et al., 2014; Yoon et al., 2011).

### Mice

All procedures were performed under Institutional Animal Care and Use Committee (IACUC)-approved protocol at Stanford University. Wild-type FVB/NCrl (207) mice were purchased from Charles River. Male mice at seven week-old age were randomly assigned to groups of predetermined sample size. All experiments with direct comparisons were performed in parallel to minimize variability. Hedgehog agonist SAG21k was delivered by osmotic pump (Alzet) over the course of two weeks at a dose of 2mg/kg/day.

### Adeno-associated virus (AAV) production

The backbones of all AAV plasmids were based on pAAV-EF1a-Cre (Addgene, 55636) with poly(A) signal replaced with bGH. AAVs were generated in HEK 293T cells and purified by iodixanol (Optiprep, Sigma; D1556) step gradients as described (Challis et al., 2019). Virus titers were measured by determining the number of DNase I–resistant vg using qPCR using a linearized genome plasmid as a standard. Purified virus was intravenously injected into anesthetized mice at 1 × 10^11 vg per mouse through the retroorbital sinus.

### Histology

Animals were euthanized and dorsal skin was excised for RNA extraction. Mice were then perfused with PBS and 4% paraformaldehyde (PFA) in PBS, and tongues and dorsal skin were post-fixed in 4% PFA for 24 hours. Tongues were processed for *in situ* hybridization according to RNAScope multiplex fluorescence kit (ACD systems) using mouse Gli1 probe (311001), followed by immunostaining as described (Lu et al., 2018). Immumofluorescence imaging was performed on laser scanning confocal microscopes (Zeiss LSM 800). Skin was processed for standard H&E staining by Animal Histology Service at Stanford University.

### RNA extraction and qRT-PCR

Skin samples were homogenized and extracted for RNA using TRIzol, followed by RNeasy Mini Kit (QIAGEN) and DNase Set (QIAGEN). Gli1 and Hprt levels were determined by one-step quantitative reverse transcriptase PCR (qRT–PCR) on an ABI 7900HT instrument using SuperScript III Platinum One-Step System with TaqMan Gene Expression Assays (Gli1, Mm00494654_m1; Hprt, Mm00446968_m1; Thermo Fisher). Normalized expression levels relative to control group were compared using Brown-Forsythe and Welch ANOVA tests with Dunnett’s multiple comparison correction.

---

**Figure S1.**
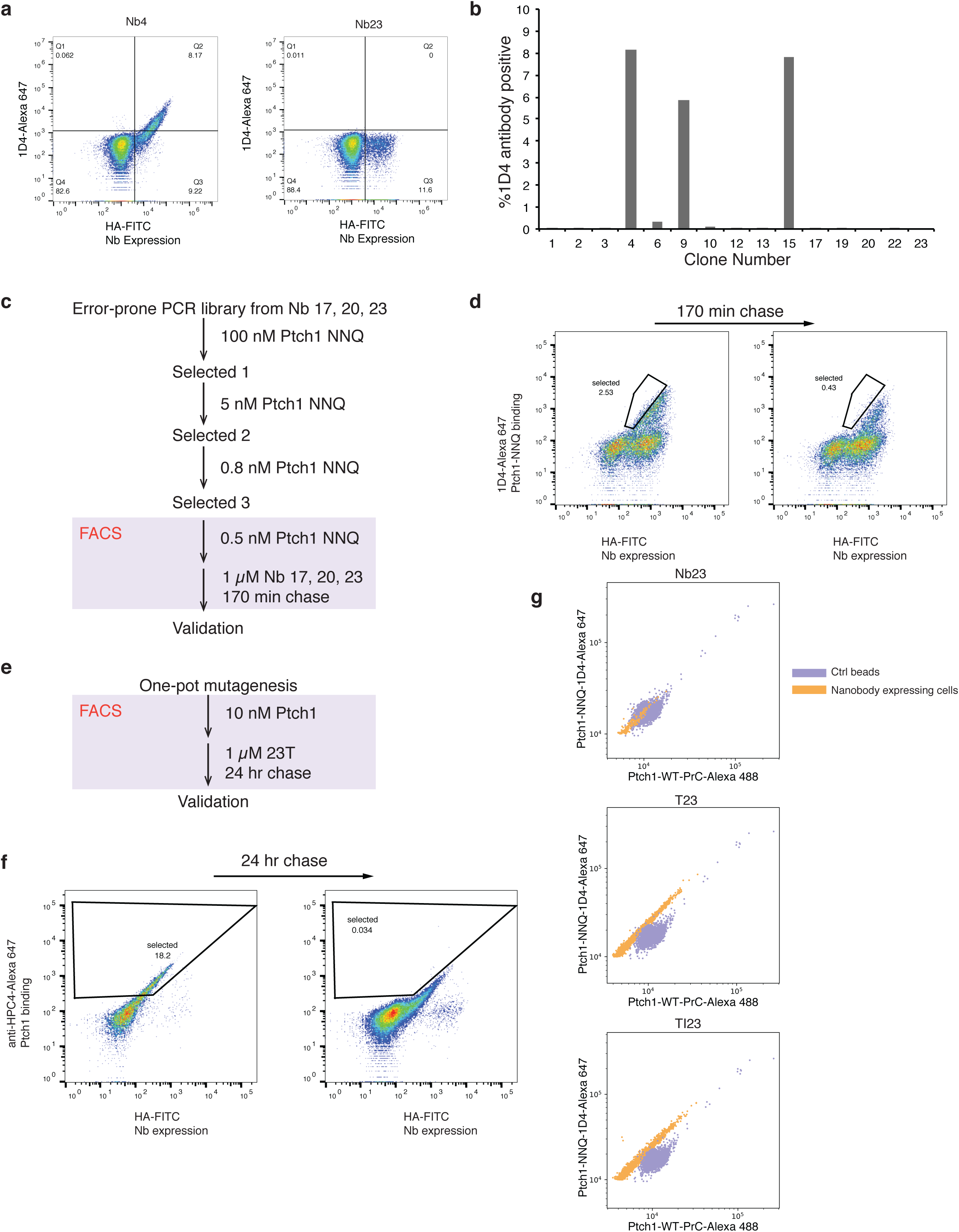
Selection of nanobody. (a) Yeast cells expressing the initial clones were stained with the antibody used during FACS to ensure that the nanobody binds directly to PTCH1 protein. As summarized in b, Clones 4, 9 and 15 showed strong binding to the antibody and are thus false-positive clones during the selection. (c) Flow chart of the first round of affinity maturation. Nanobody sequences from clone 17, 20 and 23 were mutagenized with error-prone PCR and transformed into yeast. After enriching for PTCH1 binding clones with MACS, the yeast cells are selected in FACS. In the final FACS steps, the cells were first incubated with PTCH1 to allow the nanobodies to bind and after wash, the cells were incubated with the parent nanobody proteins, to compete PTCH1 off the cell surface. FACS plots before and after the competitive chase are shown in d. The cells that retain binding to PTCH1 were selected in FACS. (e) Flow chart of the second round of affinity maturation. The sequence was mutagenized with one-pot mutagenesis and transformed into yeast. Yeast cells expressing the nanobody were selected in FACS with a similar competitive chase. The FACS plots before and after the competition were shown in f. (g) The amino acid sequences of the round 2 affinity maturation library were determined with miSeq and plotted here. The selection enriched for T77N and Y102I variants.

**Figure S2.**
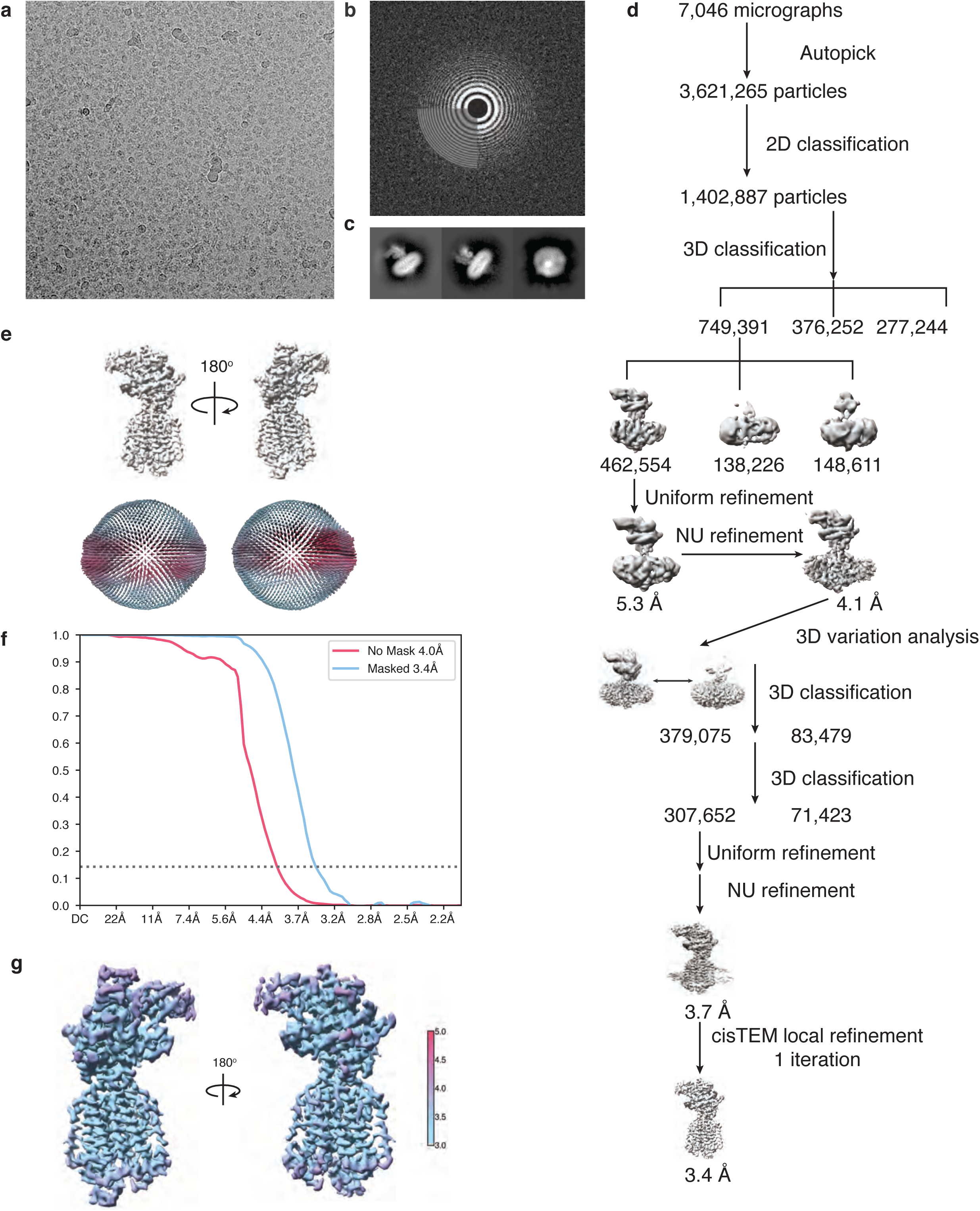
Cryo-EM data validation. (a) Protein particles are clearly visible in raw cryo-EM micrographs. (b) The parameters for contrast transfer function (CTF) are well fitted for this dataset. (c) 2D classification revealed clear views of PTCH1-TI23 complex. (d) Cryo-EM data processing was summarized in the flow chart here. All steps were carried out in cryoSPARC, except for the last local refinement step, which was performed with cisTEM. (e) The orientation of the particles is summarized in the spherical histogram here. Most particles are oriented along the equator of the protein. (f) The FSC curves of the final refinement were plotted here. The resolution of the final map is estimated to be 3.4 Å according to the 0.143 gold standard FSC. (g) Local resolution of the final reconstruction was estimated in cryoSPARC and shown in the 3D models here. Most regions were well resolved except for part of the nanobody.

**Figure S3.**
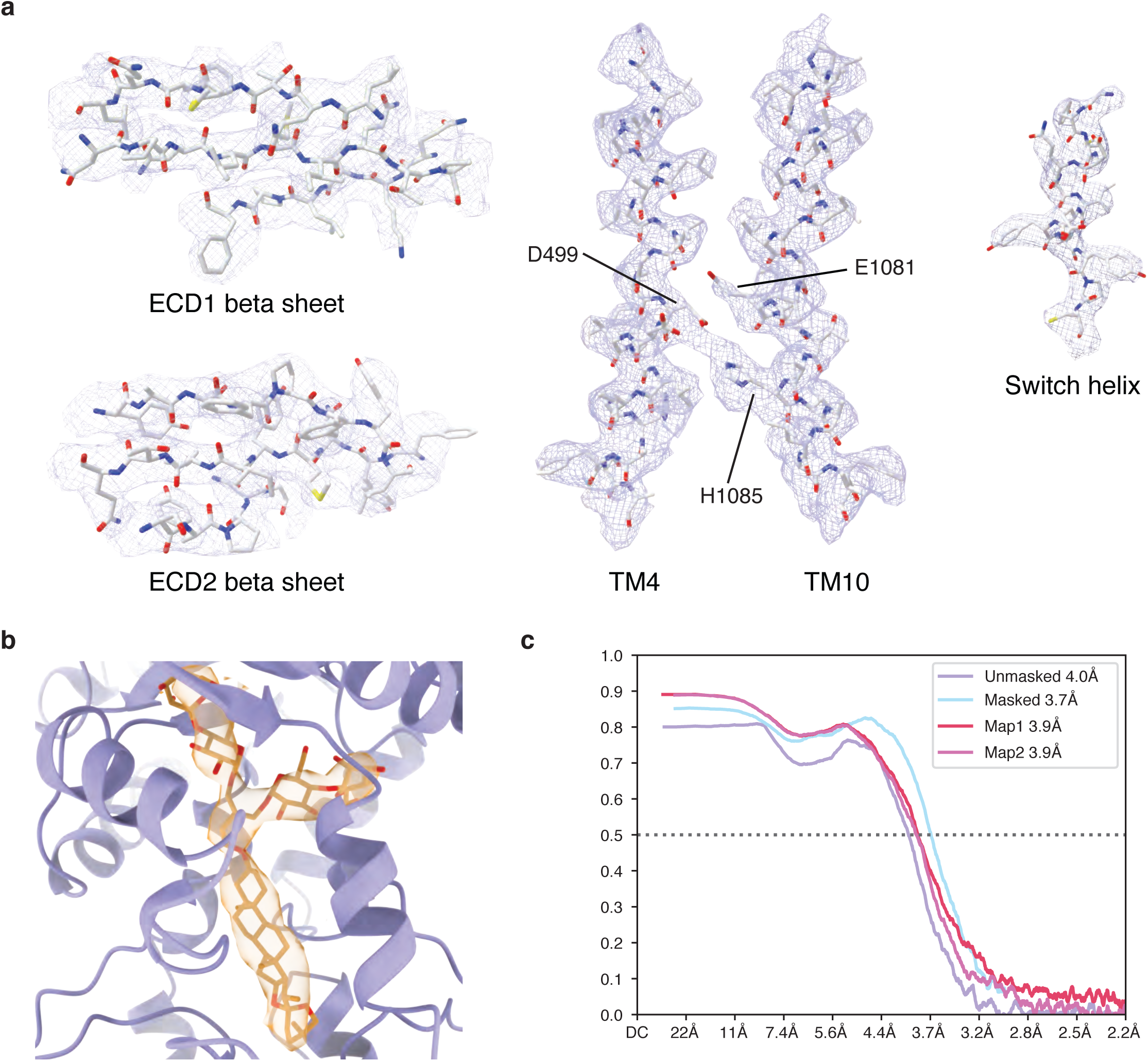
Features of the protein model. (a) The protein model fits the cryo-EM well. The high quality map enables confident modeling of not only alpha helical structures but also beta strands in the extracellular domain. Presence of clear side chain densities in the key transmembrane helices 4 and 10 enables modeling of the interaction of the key charged triad. (b) A large density present in the extracellular domain fits well with GDN and is thus likely to be a bound GDN molecule. (c) The model fits well with the cryo-EM map, as indicated by the model-map FSC curves.

**Figure S4.**
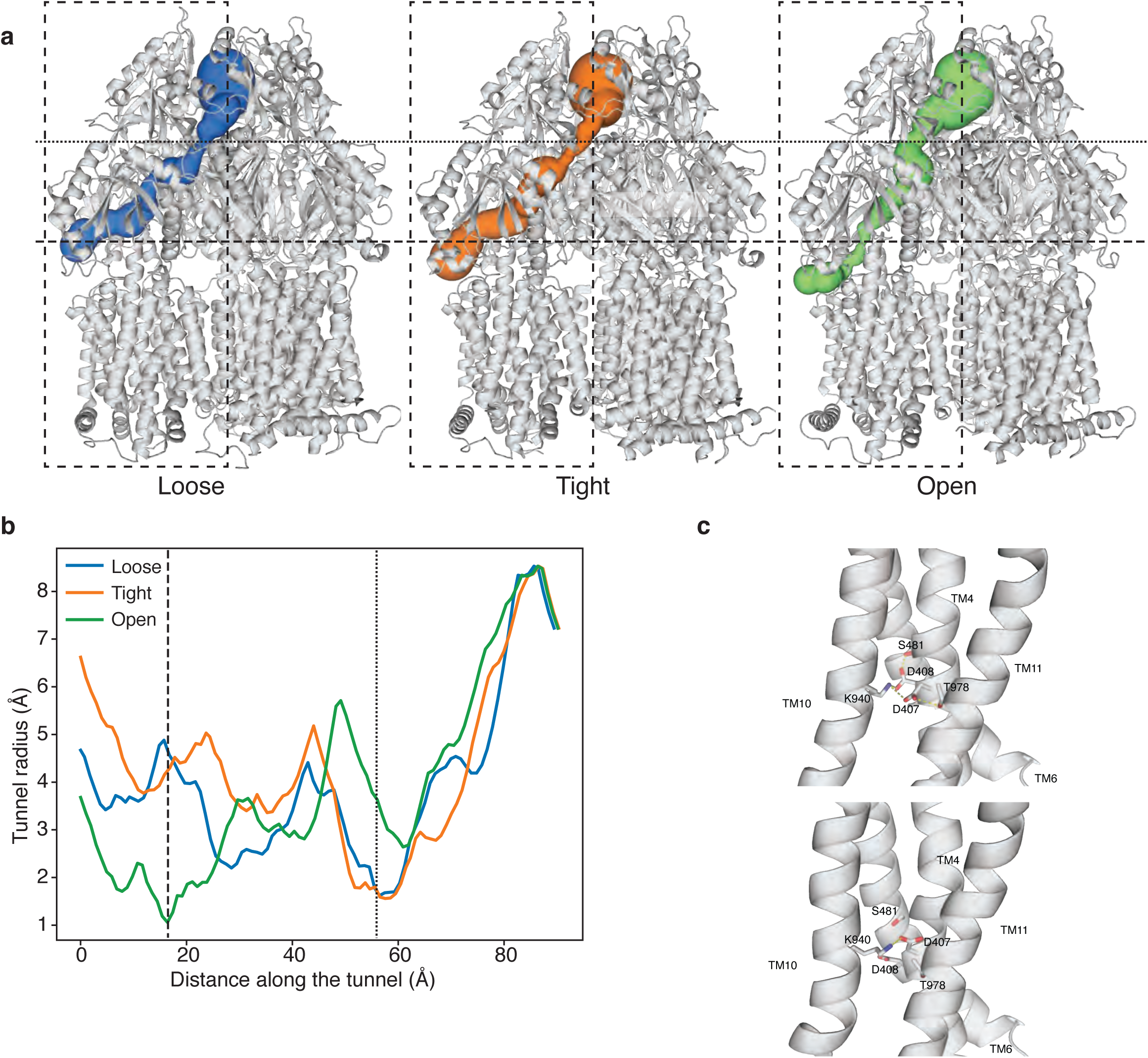
Conformational change in AcrB. (a) The transport conduit in AcrB undergoes concerted constriction and expansion in different conformations. Two narrowing parts change significantly in different conformations, and are likely to promote directional movement of substrates within the cavity. The radii of the conduit are plotted in b. (c) The interactions between the charged triad and the surrounding residues change in different AcrB conformations, coupling the transport of protons to conformational changes in the extracellular domains.

